# A real-time search strategy for finding urban disease vector infestations

**DOI:** 10.1101/2020.01.20.911974

**Authors:** Erica Billig Rose, Jason A. Roy, Ricardo Castillo-Neyra, Michelle E. Ross, Carlos Condori-Pino, Jennifer K. Peterson, Cesar Naquira-Velarde, Michael Z. Levy

## Abstract

Containing domestic vector infestation requires the ability to swiftly locate and treat infested homes. In urban settings where vectors are heterogeneously distributed throughout a dense housing matrix, the task of locating infestations can be challenging. Here, we present a novel stochastic compartmental model developed to help locate infested homes in urban areas. We designed the model using infestation data for the Chagas disease vector species *Triatoma infestans* in Arequipa, Peru. Our approach incorporates disease vector counts at each observed house, and the vector’s complex spatial dispersal dynamics. We used a Bayesian method to augment the observed data, estimate the insect population growth and dispersal parameters, and determine posterior infestation probabilities of households. We investigated the properties of the model through simulation studies, followed by field testing in Arequipa. Simulation studies showed the model to be accurate in its estimates of two parameters of interest: the growth rate of a domestic triatomine bug colony and the probability of a triatomine bug successfully invading a new home after dispersing from an infested home. When testing the model in the field, data collection using model estimates was hindered by low household participation rates, which severely limited the algorithm and in turn, the model’s predictive power. While future optimization efforts must improve the model’s capabilities when household participation is low, our approach is nonetheless an important step toward integrating data with predictive modeling to carry out evidence-based vector surveillance in cities.

## Introduction

Controlling the spread of disease vectors in real time requires locating and treating infested households before the vectors infest other houses. Vector-borne epidemics increasingly occur in cities worldwide, and detecting populations of disease vectors in large urban environments is especially complex [1, 2]. In order to manage factors that are unique to a metropolis such as contiguous and dense housing, patchy vector distribution, and urban sprawl, vector detection methods must be tailored for cities.

In the city of Arequipa, Peru, the ministry of health is nearing the end of a campaign to eliminate *Triatoma infestans*, the local vector species of *Trypanosoma cruzi*. The parasite is the etiological agent of Chagas disease, a chronic condition that can lead to serious gastrointestinal and cardiac complications. In the *T. infestans* elimination campaign, over 70,000 households were treated with insecticide during the ‘attack’ phase of this effort, and are now in a community-based surveillance phase. In this phase, residents are encouraged to report suspected *T. infestans* infestations, which are then verified by a trained entomological inspector. If *T. infestans* are found in or around the home, the area will be treated with insecticide, along with all neighboring homes. To complement the community based efforts, trained vector control personnel also proactively inspect a proportion of houses in each district for vector infestation.

As a result of these two ongoing surveillance actitivies in Arequipa, new infestation data for the city are constantly being collected. However, the data are not used to inform subsequent surveillance activities. Therefore, we aimed to develop a tool for entomological inspectors that estimated household infestation risk based on data. We also wanted the tool to incorporate new infestation data as infested houses are found and treated, in order to predict the locations of other infestations in close to real time. Here, we present the model we developed to proactively search a landscape for infested houses, folllowed by model simulation results and some preliminary field testing outcomes in Arequipa.

## Methods

Our models build on a rich literature of susceptible-infectious-removed (SIR) models. Likelihood-based methods have been developed to fit SIR models to data when the infection times and total epidemic size are known [3–5]. These methods were extended, using a reversible-jump Markov Chain Monte Carlo algorithm (RJMCMC), to cases when these parameters are unobserved [6–8]. Later, Jewell et al. extended the methodology to notifiable diseases by describing the length of time from the unobserved infection time to some later, observed ‘notification’ time [9,10]. For our model, we further extended these methods by applying them to both cross-sectionally and longitudinally collected data on vector populations. By adding structured growth of the vector populations, we were able to estimate the ‘infectious’ period of infested household (i.e., the time whenvectors disperse from the infested home to susceptible homes), and thereby fit an SIR model to the data. Our model incorporates both spatial heterogeneity and the severity of infestation into the transmission process.

Our SIR model is set to the individual household level. We use a stochastic epidemic modeling approach to estimate the posterior probabilities of infestation of each house within a given region. At any given time point, a house may be in one of the following states:

1. **Currently infested and not yet treated:** We assume that these houses continue to be infested until treatment.
2. **Previously infested and treated within the last 100 days**: We assume these houses are removed from the system, and not susceptible to re-infestation.
3. **Previously infested and treated more than 100 days ago**: We assume these houses are susceptible, and may be infested any time after the effective interval of insecticide, thought to be 100 days post-treatment [11].
4. **Previously inspected and known to have been free of vector infestation at some point in the past**: We assume these houses are susceptible at any point after the previous inspection.
5. **Never inspected and no information ever known**: We assume these houses are susceptible at any time point since the initial infestation in the area.

For each house, we incorporate data collected from both active and passive surveillance into our approach. The vast majority of inspected houses have not been infested during the surveillance phase; we assume each house is uninfested at the time of a negative inspection, but may have been infested since.

Our model uses the number of insects found in each inspected house during the most recent inspection, in combination with a household-level insect population growth model (described later) to estimate the unobserved true date that the house was first infested. Dates of insecticide treatment (in the surveillance data used) were all observed, and treatment was assumed to be 100% effective.

### Notation

Before introducing the model and likelihood, we define some notation:

- Define *I_i_*, *D_i_*, and *R_i_* as the infestation time (date on which it was first infested), detection time (date the infestation was detected), and treatment time (i.e., date) of the *i*th house, respectively.
- Define *h_ij_*(*t*) as the probability that house *i* infests house *j* at time *t*.
- Define *b_i_*(*t*) as the number of insects in the *i*th house at time *t*, and *b_i_* is the set of all insect counts of house *i* over all time points.
- Define *r* as the growth rate of the insect population within the house. Due to the slow moving nature of household re-infestations, we describe the growth rate as the insect population increase per time unit *t* of 90 days. We may estimate *r* from the data if there are enough observed infestations. Otherwise, we fix *r* to values identified from previous studies [12].
- Define *K* as the carrying capacity, or the number of insects a single household can support. We assume each house to have the same carrying capacity, and pick *K* to be large enough that it is a reasonable estimate of a carrying capacity seen in Arequipa.
- Define λ_*t*_ as the expected number of insects at time *t* given the time of infestation *I_i_* according to the assumed population growth model (we assume one insect at the time of infestation).
- Define *N*, *N_I_*, and *N_D_* as the total number of houses, number of infested houses, and number of detected infested houses, respectively, at the current time (today), *T*_max_. *N* and *N_D_* are observed data. However, *N_I_* is unobserved, and is presumably larger than *N_D_*.
- Define *β* as the probability of successful invasion given a migrating insect. We estimate *β* from the data.
- Define *t*_insp,*i*_ as the inspection time of the *i*th house, and *T*_max_ as the current time

### Likelihood

Next, we introduce the complete-data likelihood (the likelihood if we observe infection status and insect counts at all houses at all times), followed by model descriptions for each part of the likelihood. We will then describe prior distributions and inference algorithms.

The complete-data likelihood is:

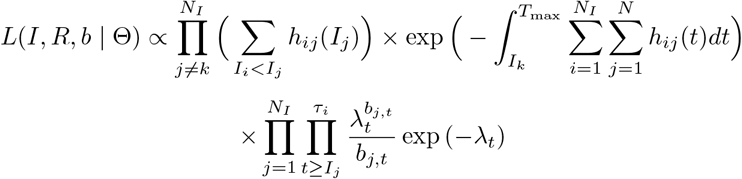

where Θ = {*β*, *r*, *K*, λ_*t*_, *I_κ_*}, *τ_i_* = min(*R_i_*, *T*_max_) and *I_κ_* is the initial infestation time. The parameters *β* and *r* are unknown, and therefore will need to be assigned prior distributions. We fix *K* and *I_κ_* to plausible field estimates. If we do not have many infestated homes in a region of the city, we may also fix *r*, but in our simulations we demonstrate that our model is capable of estimating both *β* and *r*. In our model, *R_i_*, *T*_max_, *N_D_*, *N* are observed and *N_I_* and *I_j_* is unobserved. We observe *b*_*i*, *t*_insp__, but all other *b_i_*(*t*) are unobserved, and will be imputed using a household-level insect population growth model. λ_*t*_ is also estimated using the population growth model; details of the estimation procedure are described later.

The first piece of the likelihood describes the probability that the *i*th house infests the *j*th house at the time the *j*th house was infested, *I_j_*. The second piece of this likelihood describes the cumulative infectious pressure. The infectious pressure captures the effect over time of each infested house on every other uninfested house. In other words, at a given time *t*, a house that is surrounded by infested houses will be much less likely to escape infestation than a house that is surrounded by insect-free houses. This kind of pressure, over time, is captured by the second term in the likelihood. The third piece of the likelihood describes the probability of observing the number of insects at each time point of each infested house, which we assume follows a Poisson distribution. Specific models for the transmission process *h_ij_*(*t*), which incorporates spatial heterogeneity, and the insect count rate λ_*t*_, are described below.

It has previously been shown that the integral in the likelihood can be written in a simpler and intuitive way ([9]):

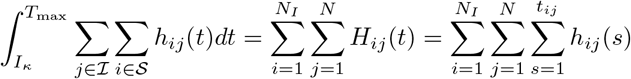

where *t_ij_* = min(*R_i_*, *I_j_*, *T*_max_) − min(*I_i_*, *I_j_*). The likelihood then becomes:

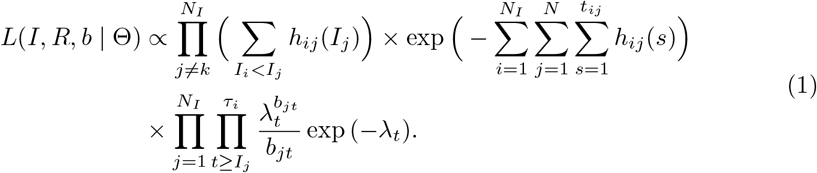

To account for the possibility that a given house *j* is infested by two houses at the same time point, we calculate the probability that house *j* is infested at time *t* by calculating 1 − *P*(house *j* avoids infestation at time *t*):

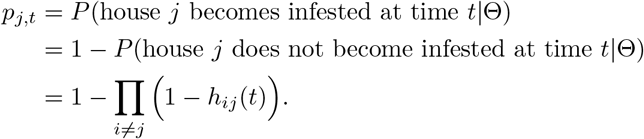

In addition, since we are working in the Bayesian framework, we put prior distributions on the parameters that we are estimating. We chose a beta prior distribution on *β* and a gamma prior distribution on *r*.

### Insect infectiousness

We allow for heterogeneous transmission relative to the number of insects in each infested house. We hypothesize that houses with more vectors are more likely to infest their neighbors than houses with only a few vectors. We characterize this heterogeneity in infectivity using a house-level population growth model. We use the Beverton-Holt model, which has two parameters: a house-level carrying capacity and insect growth rate [13,14]. The Beverton-Holt model, shown in Fig 3, is a key element of our extension that captures the distribution of the infectious period.

The model is:

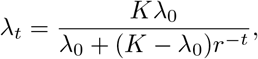

where λ_0_ is the number of insects at the initial time, λ_*t*_ is the expected number of insects at time *t*, *K* is the carrying capacity, and *r* is the growth rate per generation. For Chagas disease in Arequipa, we assume that λ_0_ = 1 and *K* = 1000, implying that each household infestation begins with one insect, and has a carrying capacity of 1000 insects. In truth, the carrying capacity is unknown, and we have conducted sensitivity analyses on this assumption (in Supporting information). However, we believe this is a realistic estimate of carrying capacity based on extensive experience working in the field in Arequipa. We estimate *r* through the RJMCMC algorithm.

Using this model, when there are an expected λ_*t*_ insects in a given house, the time since the population was one insect can be solved for explicitly:

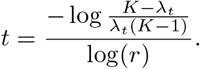

We assume that the rate at which insects migrate to find new houses from any given infested house is the difference between the unbounded and bounded growth rates. As described above, the expected insect population size within the infested house from which the bugs spread at time *t* under the bounded (Beverton-Holt) model is λ_*t*_. The bounded growth rate is therefore 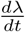. If there were unlimited resources, then insect populations would grow exponentially. Define 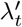 as the number of insects expected at time *t* assuming exponential growth, 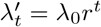. The unbounded growth rate is 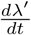. The ‘infectiousness’ of each house, meaning the rate at which vectors from the infested house spread to other houses, is the difference in slope between these two models at a given time point *t*.

In practice, we are unable to observe the insect count within a given house at every time point. At each inspected house, we observe the insect count (i.e.,number of insects found by the inspector) at one time point, and the rest can be imputed using data augmentation. We assume insect counts follow a Poisson distribution, centered around the Beverton-Holt function, λ_*t*−*I_i_*_ where *t* − *I_i_* is the time since infestation:

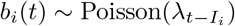

where *b_i_*(*t*) is the number of insects in house i at time *t*. This Poisson distribution is the third component in the likelihood (1).

We can now incorporate information about insect counts, spatial heterogeneity, and house infectiousness into the vector transmission function. We assume the following vector transmission function *h_ij_*(*t*):

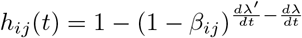

where *β_ij_* = *β* × *δ_ij_*. We estimate *β* through the RJMCMC, and *δ_ij_* is the estimated spatial kernel, described below.

Although this transmission function is mathematically complex, it can be interpreted more simply, as follows: Given the infestation status of house *i* and the distance between houses *i* and *j*, *β_ij_* describes the probability that house *i* infests house *j*. Thus, 1 − *β_ij_* describes the probability that house *j* escapes infestation from house *i*. The probability that house *j* escapes infection exponentially decreases as the number of insects in house *i* increases. We characterize the dynamic of migrating insects as the difference in population growth between unbounded population growth and bounded population growth. Thus, subtracting this quantity from one gives the probability that house *i* does in fact infest house *j*.

The transmission function appears twice in likelihood (1), both as the infectious pressure of each infested *i* house on every *j* house, 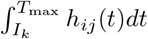 and cumulative hazard function of each infested house *i* on each *j* house, 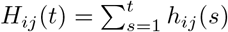.

### Spatial dynamics

Since *T. infestans* migration is heterogeneous, we incorporate this spatial heterogeneity into the hazard function, *h_ij_*(*t*) [15]. It is important to note that any number of spatial kernels can be implemented into this approach. Barbu et al [16] identified a modified exponential transmission kernel for *T. infestans* migration that incorporates both distance between houses and a city block indicator. We use this kernel to describe the probability of an insect migrating from house *i* to house *j* given the distance *d_ij_* between the two houses. Using this kernel, we also incorporate city streets as barriers, since previous studies have shown that *T. infestans* are less likely to move a given distance between city blocks compared to within city blocks [16,17]. We use normal prior distributions on *ζ* and *ρ*, centered around values identified previously [16]:

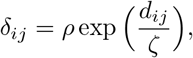

where *d_ij_* defines the Euclidean distance between houses *i* and *j*, *ζ* ~ N(9.00, *σ_ζ_*), and *ρ* = 1 if houses *i* and *j* are on the same city block. If houses *i* and *j* are not on the same city block, *ρ* ~ N(0.30, *σ_ρ_*).

### Model fitting

To fit our model, we wrote a reversible-jump Markov Chain Monte Carlo (RJMCMC) algorithm using R software v3.2, and conducted likelihood calculations using the ‘Rcpp’ package [18]. The RJMCMC algorithm allows for changes in the dimensionality of the parameter space [6,19]. We allow for the possibilities of ‘adding’ a potentially unobserved infestation, removing an already added unobserved infestation, or shifting an infestation in time. Although the parameter estimates converge quickly, we run the RJMCMC for a minimum of 1 million iterations, to ensure adequate convergence of the posterior probabilities of infestation, which takes about 12 hours. After running the algorithm for *M* iterations, each house with an unknown infestation status is added as infested for *m_i_* ≤ *M* iterations. Thus, the calculated posterior probability of infestation of each house is 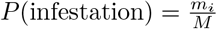. For more details on the algorithm and convergence, see Supporting information.

### Simulations

We tested our algorithm on simulated data generated on a real landscape. We chose a subset of the study region in Arequipa (173 houses) in which to simulate an epidemic and then we randomly selected one house to be the initial infestation, *κ*. We set this house to have 1 insect at time *t* = 1, *b*_*κ*, 1_ = 1. We then carried out forward simulations using the Beverton-Holt model *b*_*κ*, *t*_ ~ Poisson(λ_*t*_) until *t*_max_. At time *t* = 2, house *κ* infests each other house *j*, *j* ≠ *κ* with probability *h_κj_*(*t*) = *h_κj_*(2). As more houses become infested, these houses can then infest other houses. If house *i* is not yet infected at time *t*, then *h_ij_*(*t*) = 0 (ie. the probability that house *i* infests house *j* at time *t* is zero). The inspection time of each house was randomly selected *t*_insp, *i*_ ~ Uniform(*I_i_*, *t*_max_). The initial infestation time *I_κ_* is the only infestation time that is considered observed. To simulate unreported infestations, we randomly chose 1/3 of the infestations to treat as unobserved.

We simulated the data under three sets of parameter values {*β*, *r*}={0.3, 2.0}, {0.7, 2.5}, {0.05, 3.0}. To create realistic data, we were limited to simulated datasets under a subset of parameter regimes in which vector spread occurs. For each simulation, a wide beta prior distribution is used for *β* and a wide gamma distribution is used for *r*, both centered around the true value.

Under each set of parameter regimes, we simulated 200 datasets. For each dataset, we ran 3 chains at different starting values. We ran each chain for 500,000 iterations, and discarded the first 5000 iterations as burn-in. We assessed convergence by plotting the rankings of all chain pairs (more details in Supporting information). From each simulation, we calculated the sensitivity and specificity of the model by recovering occult infestations under various thresholds of inspection criteria. We created receiver operating characteristic (ROC) curves to examine the simulation results (Fig 1). We also estimated the parameters of interest, *β* and *r*. For each simulated dataset, we recorded the median,the 95% credible interval and area under the curve (AUC), and the mean.

**Fig 1.**
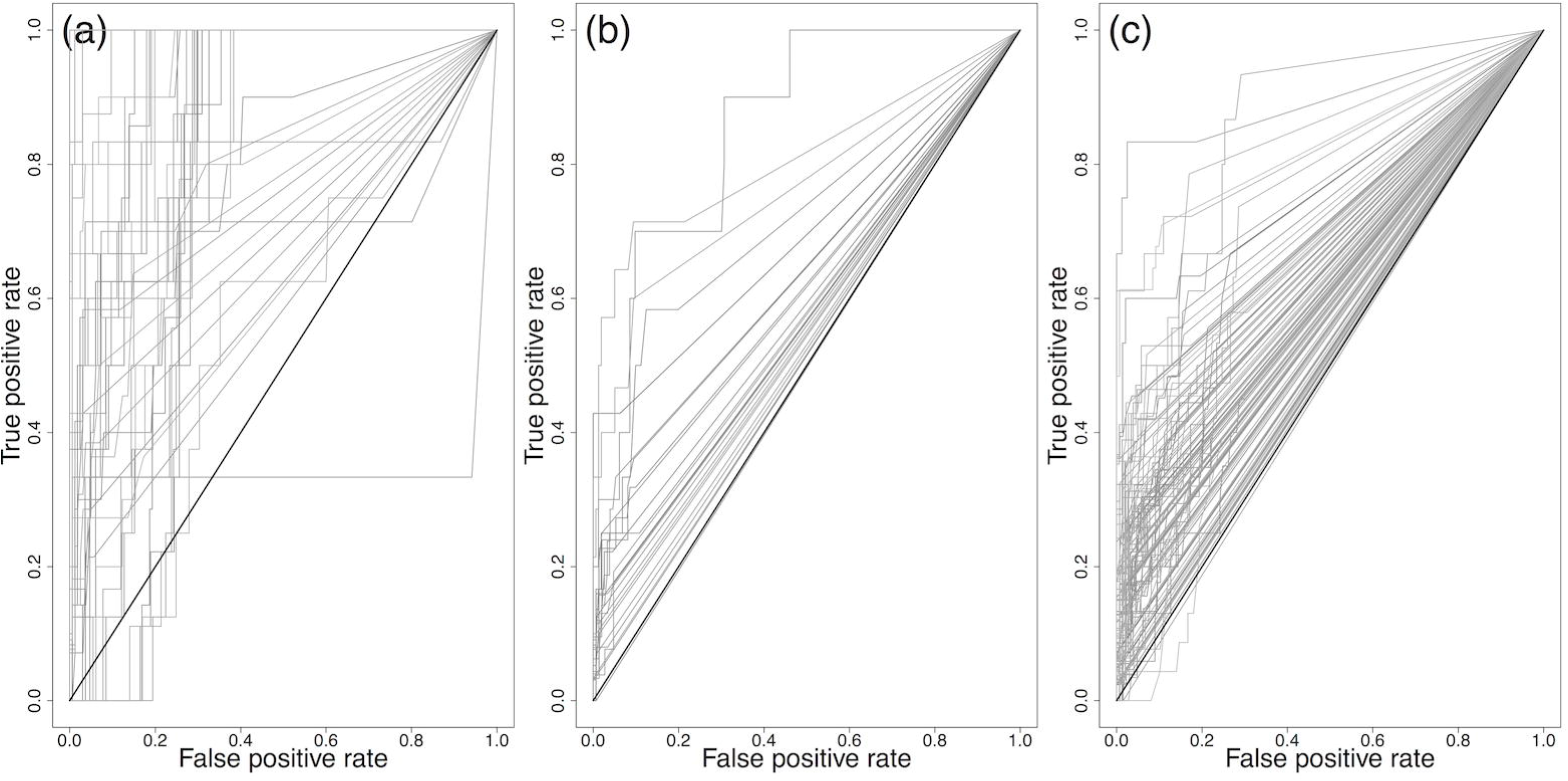
ROC curves describing the ability of the RJMCMC to uncover unreported infestations in a simulated vector-borne epidemic. In each simulation, 1/3 of infestations were randomly removed to recover. 50 randomly selected ROC curves of each parameter regime are shown. (a) {*β*, *r*} = {0.02,1.6} (b) {*β*, *r*} = {0.7,1.8} (c) {*β*, *r*} = {0.3, 2.1}

### Preliminary field testing

Between October and December of 2015, we pilot tested our model in Arequipa in near real-time. The pilot took place in three adjacent communities, each containing between 551 and 669 houses. Each community had been treated with insecticide in 2004; we used data collected during treatment to calculate prior probabilities of infestation for each household, following the methods of Barbu [20]. Since 2004, residents have reported suspected infestations, which were then verified by our field staff and Minstry of Health entomological inspectors. When suspected infestations were confirmed, houses adjacent to the infested home were inspected as well. Thus, since 2004, some houses have been inspected again at unique time points, either due to a resident report or an infested neighboring house. The number of observed infestations in the three study communities between 2004-2014 was 9,10, and 11. Observed insect counts in the infested houses during this time period ranged from 0 to 212, with the median insect count being 3. Due to the low number of observed infestations and insect counts in the communities, we did not attempt to estimate the insect growth rate, *r*. Instead, we drew *r* from a normal distribution centered around 3.66, as found in [12], and we just estimated *β*. The posterior distribution of *β* at the beginning of the study is summarized in Fig 2. The *β* estimates were similar between communities. We observed a wide 95% credible interval on the posterior distribution of *β* across the three communities. The estimates of *β* remained consistent throughout the pilot study.

**Fig 2.**
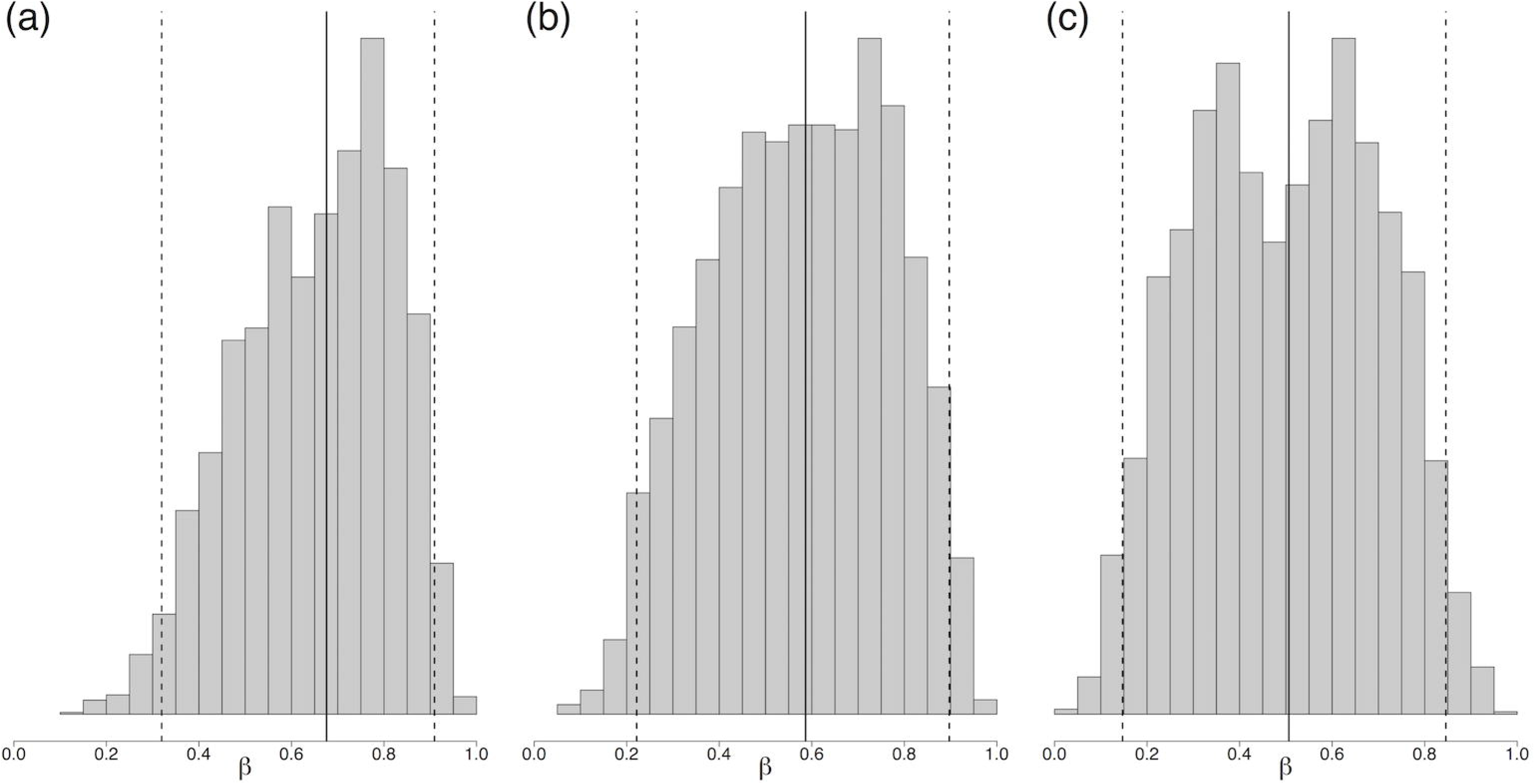
Posterior estimate of *β*, the probability of successful invasion of a migrating insect, at the beginning of the inspection campaign in each community, indicating median and 95% credible interval of RJMCMC chains: (a) Community 1: 0.67 (0.33, 0.91) (b) Community 2: 0.59 (0.22, 0.90) (c) Community 3: 0.51 (0.15, 0.85)

**Fig 3.**
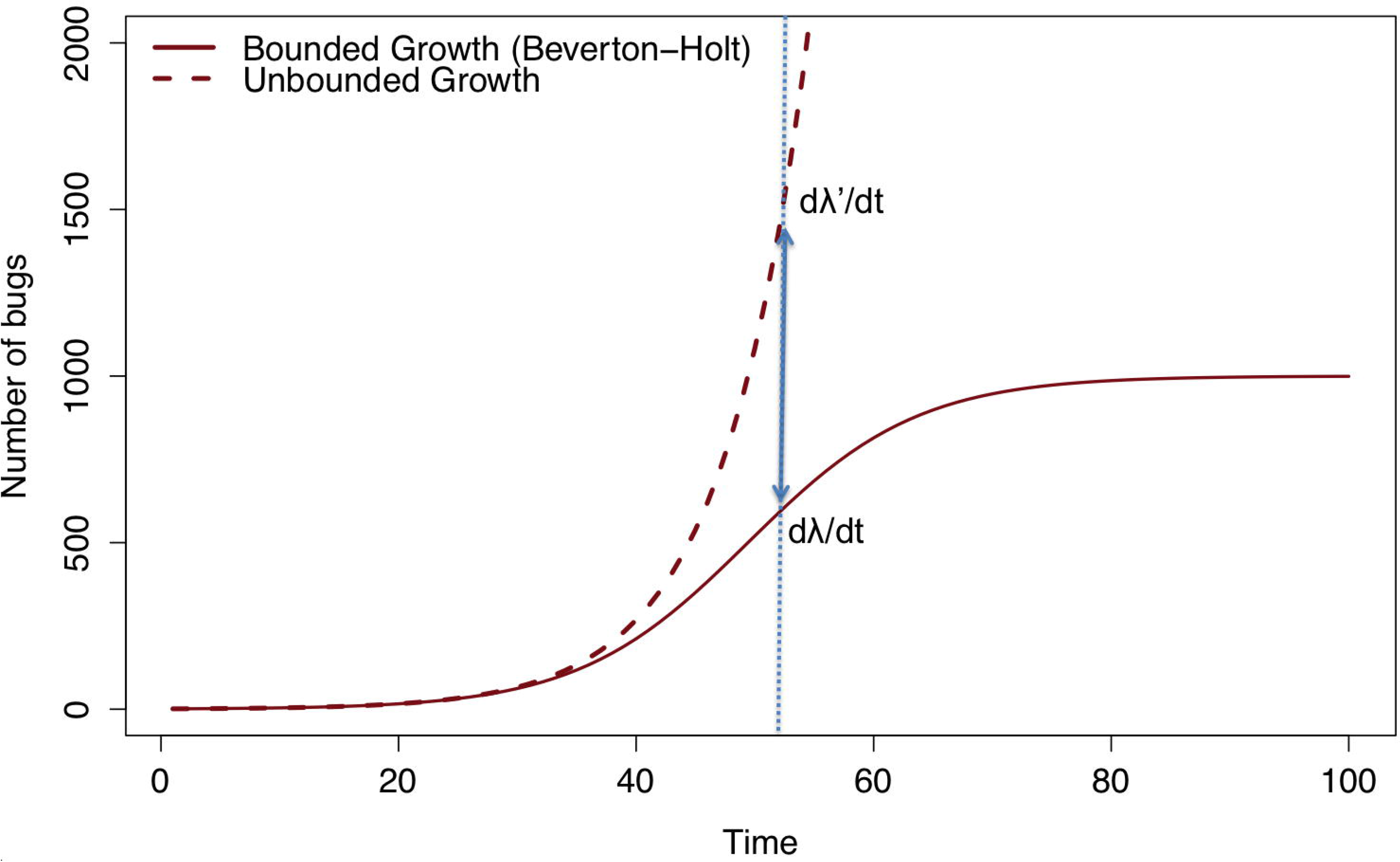
Bounded growth using the Beverton-Holt model compared to unbounded growth. Difference in slopes at time of new infestation was incorporated into hazard function to quantify infestation severity into the probability of infesting a neighboring house.

**Fig 4.**
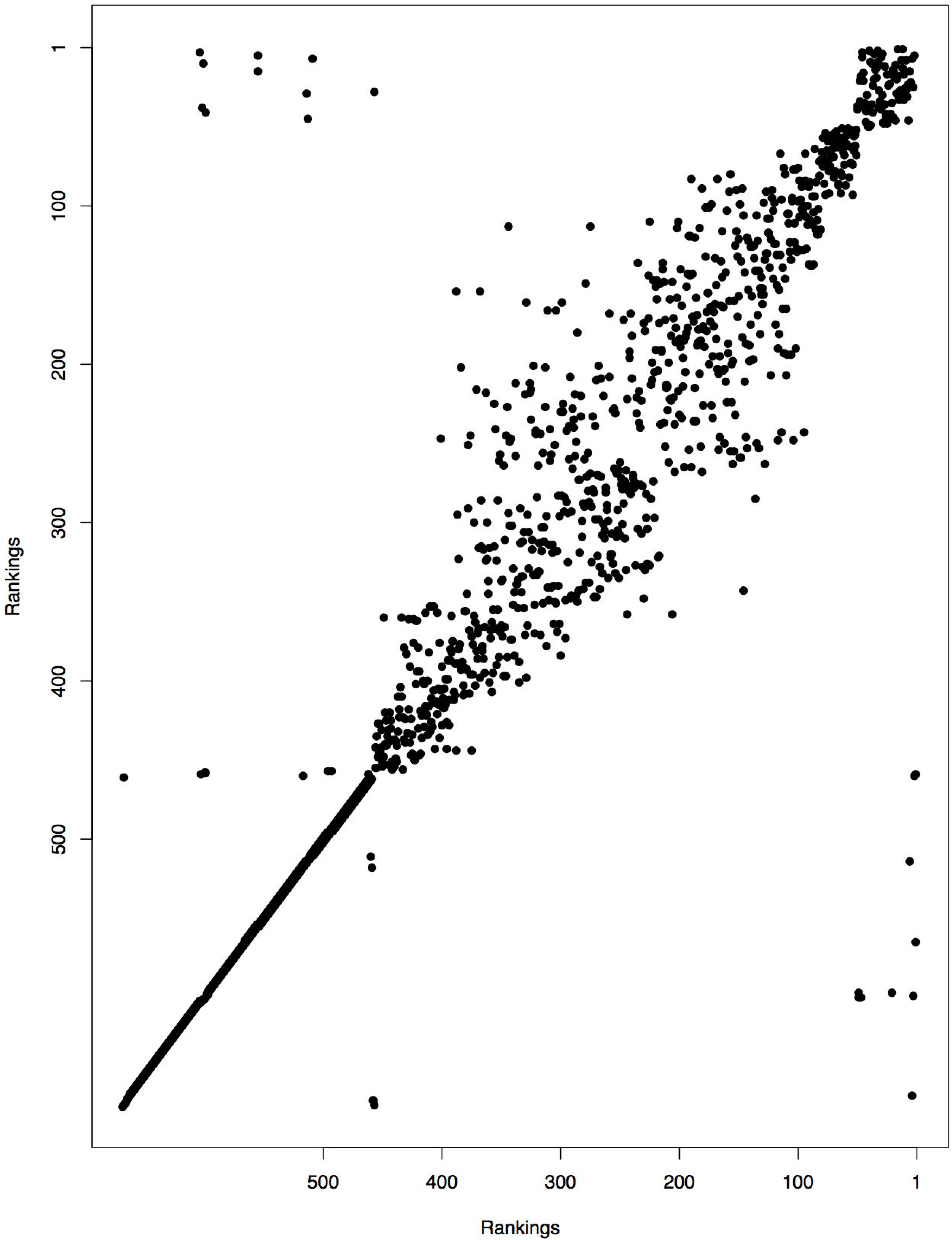
Ranking of each house across 3 RJMCMC chains. There were a few houses that changed significantly, but most houses remained within a few rankings between chains. In cases of discrepancies, the median ranking across the chains was used to determine which houses to search.

**Fig 5.**
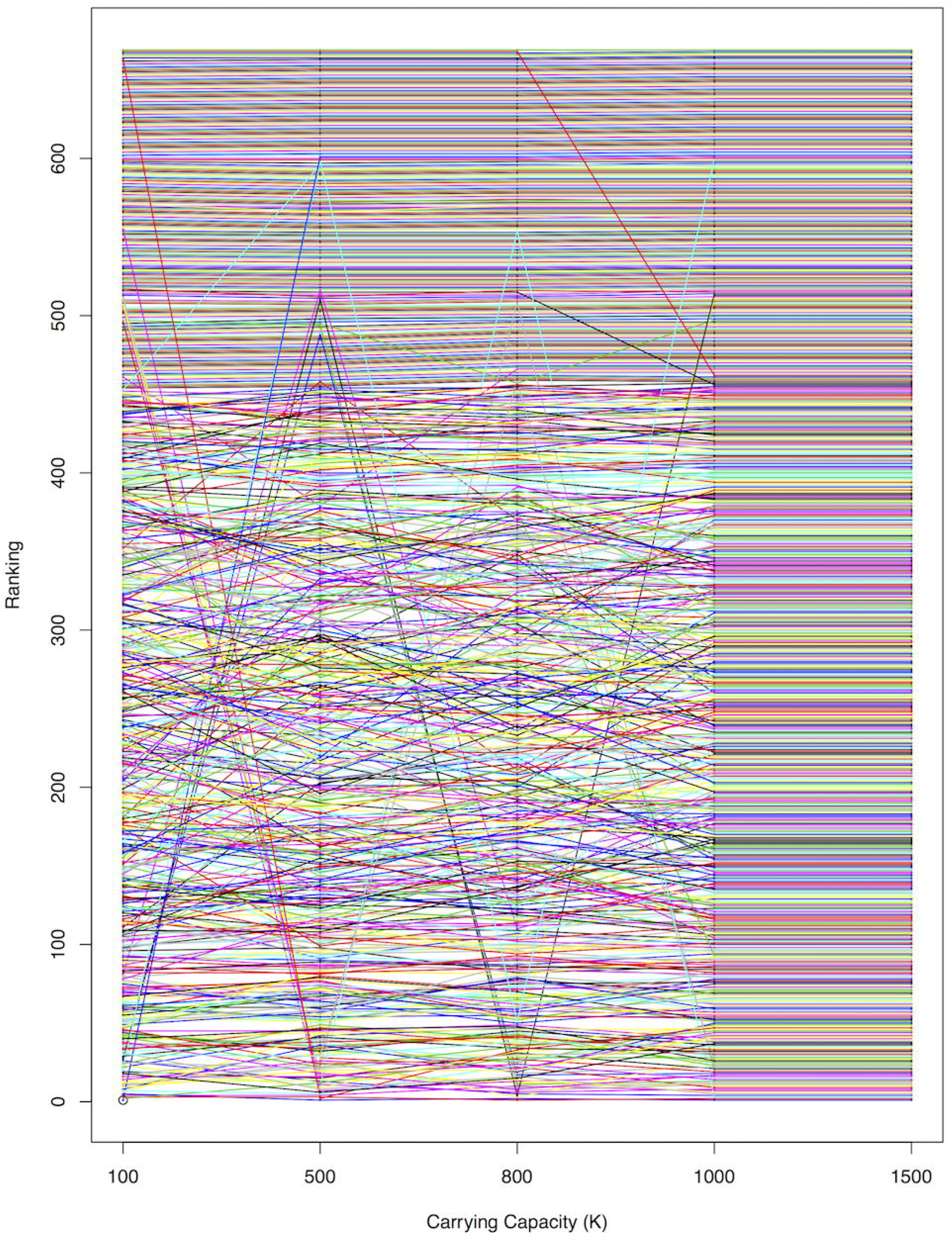
Ranking of each house across 5 potential carrying capacities. There were a few houses that changed significantly, but most houses remained within a few rankings between carrying capacity values.

In the field, each day inspectors searched houses with the highest posterior probability of infestation as predicted by the model. Each inspector traveled only within one community per day. After each day of inspections, we ran five chains, and obtained a ranking for each house by using the median ranking across the chains for each locality. We ran the model for as many iterations as possible, which was over a million iterations, in order to include observed data from the most recent day and to create the new ranking list for the following day. Houses in which the resident refused inspection or did not answer the door were kept in the algorithm as unknown infestations; those in which the resident did not answer the door may be revisited at a later date, while those in which inspections were refused were removed from the pool. Abandoned houses were assumed to be uninfested, and were not included in the algorithm. Houses outside of the study sites but within 50 meters of the study site border were included in the model, but not in the potential inspection pool. Including these proximal houses allowed for the possibility of vectors entering the study site from a neighboring locality.

## Results

### Simulations

In each simulation, our median estimates of *β* and *r* were close to their true values (Table 1). The AUC suggested that the model performs best under low values of *β*, i.e., when insects dispersing from a given house have a low probability of successfully invading other houses. The acceptance rate of the RJMCMC ranged from 20-50%. These results suggest that our approach accurately estimated the parameters of interest.

**Table 1.** Simulation results. Each estimate represents the mean of given quartile for the set of 200 replicates. We use the median estimate to verify our approach obtains accurate parameter estimates and report the median AUC of each parameter set.

| Parameter Estimation |  |  |  |  |  |  |  |
| --- | --- | --- | --- | --- | --- | --- | --- |
| | $\hat{\beta}$ | | | $\hat{r}$ | | | AUC |
|  | 2.5% | 50% | 97.5% | 2.5% | 50% | 97.5% |  |
| $\beta = 0.02, r = 1.60$ | 0.0004 | 0.03 | 0.60 | 1.27 | 1.62 | 1.94 | 0.83 |
| $\beta = 0.70, r = 1.80$ | 0.31 | 0.76 | 0.98 | 1.21 | 1.73 | 1.95 | 0.57 |
| $\beta = 0.30, r = 2.10$ | 0.05 | 0.26 | 0.62 | 1.94 | 2.04 | 2.13 | 0.61 |

### Pilot test

We made a total of 835 visits to 409 distinct households that were predicted by the model to be at high risk of infestation. Of these houses, 135 (33%) granted permission for inspection for *T. infestans*. The remaining homes either refused inspection, did not answer the door, or were abandoned. In the 135 homes that were inspected, no *T. infestans* were found. The absence of vector infestation in these homes indicates that the *T. infestans* infestation prevalence in the area is much lower than we had anticipated at the beginning of the study, even though inspectors were directed to areas that were close to previous reports of infestation.

## Discussion

Here, we present a novel dynamic model designed to enhance disease vector surveillance by updating posterior probabilities of infestation as inspection data are collected. Simulations based on historical data indicated that the model performed well, although its field application was limited by practical constraints, especially resident participation.

Despite these preliminary limitations, our approach has valuable aspects that could improve vector surveillance and control, especially in cases where detailed data are not available. For example, we commonly observe the infection status of a subset of individuals at some time point after the infection occurred [1, 21, 22], and the actual timepoints of each infection are often unknown. Methods have been developed to handle incomplete data from infectious disease outbreaks, including unobserved infection times and incomplete epidemics [7, 9]. Alternatives to likelihood-based methods include approximate Bayesian computation [23, 24] and synthetic likelihood [25]. Other methods of inference have been used as well, such as iterated particle filtering [26]. In our approach, we extended these methods by taking a Bayesian model developed by [10] for notifiable diseases and applying it to cross-sectional observations of insect infestations that can be updated in near real-time as new data are collected. We provided a framework that can be readily extended to other vector infestations, such as the current global bed bug epidemic.

Limitations of the model itself include its dependence on several assumptions that are made to retain a tractable likelihood. We assume a specific spatial kernel and insect population growth model, which is key to our ability to perform a likelihood-based analysis by enabling the estimation of the unobserved infestation times. We tested the sensitivity of our results to fixed parameters in the insect population growth model, and we found that the model was not very sensitive to different insect carrying capacities (see Supporting information). In addition, we assume perfect inspections and that insecticide application is one hundred percent effective. Although the inspectors are highly skilled, there is always the possibility of heterogeneity in the detection accuracy of active vector surveillance, as found in other studies [21]. It is especially difficult to detect early stage nymphs, which are small and difficult to see. Similarly, while insecticide treatment is known to be effective against *T. infestans*, we cannot verify that all prior domestic infestations were completely eliminated, especially in severe cases. Lastly, determining convergence of the posterior probabilities of infestation is complex. The chains are qualitatively consistent in their rankings of houses (in terms of posterior probability of infestation), but the rankings themselves are not identical between chains, indicating limited convergence. We used the median ranking across chains to minimize this limitation.

Our modeling approach is purely exploitative; inspectors were sent to houses with the highest posterior probability of infestation given the current knowledge of the system. An inspection algorithm that appropriately balances both an exploitative and explorative model may be more appropriate in future search strategies. Our model also requires a modicum of spatio-temporal data, including the location of infested houses. The model cannot be applied to areas in which there are no previously identified infested households, including any area that is newly entering surveillance. In these cases a more general spatial-temporal model such as that described in [17, 27] may be more appropriate, at least until sufficient information to fit the reversible jump MCMC has accumulated.

## Conclusion

The development and testing of mathematical tools for use in vector control is an important step toward integrating computational methods into epidemiological surveillance. Our approach uses a dynamic model that incorporates ecological and infectious disease principles to guide real-time field investigations, enabling inspectors to make informed vector surveillance decisions. While field testing revealed key areas for improvement, the framework is still an important step towards evidence-based entomological surveillance, which is a critical component of integrated vector management.

## Acknowledgments

We gratefully acknowledge the invaluable contributions of the Ministerio de Salud del Peru (MINSA), the Dirección General de Salud de las Personas (DGSP), the Estrategia Sanitaria Nacional de Prevención y Control de Enfermedades Metaxénicas y Otras Transmitidas por Vectores (ESNPCEMOTVS), the Dirección General de Salud Ambiental (DIGESA), the Gobierno Regional de Arequipa, the Gerencia Regional de Salud de Arequipa (GRSA), and the members of the field and laboratory teams at the Zoonotic Disease Research Laboratory in Arequipa. This study was supported by National Institutes of Health grants 5T32AI007532 and 5R01AI101229.

## Supporting information

### Growth Dynamics

See Fig 4.

### Algorithm details

We update the likelihood using the following algorithm:

1. Initialize *β*^1^ and *r*^1^.
2. Initialize infection times *I*^1^. Initial infestation *I_κ_* = 1 and all other observed infestations initialized at *I* = 2. All other infestations set to *I* = ∞.
3. Initialize *t*_insp, *i*_ and *R_i_*. If house *i* has been inspected andor treated, these are set to the respective times. All dates are converted to time since initial infestation, using 90 day intervals. For example, if house *i* was infested 200 days after the initial infestation, this infestation time is considered *I_i_* = 3. If a house has not yet been inspected and treated, *t*_insp, *i*_ = *R_i_* = ∞. *T*_max_ is set to the present day.
4. Initialize insect counts of each infested house given the infestation times *I*^1^ This is done using the Beverton-Holt Model:

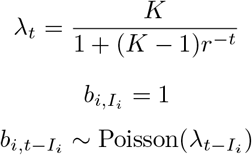

Replace *b*_*i*, *t*_insp, *i*__ = *B*_*i*, *t*_insp, *i*__, the observed insect counts.
5. Update *r*^*m*+1^. Propose *r*⋆ ~ Normal(*r^m^*, 0.05). Update all insect counts *b*⋆ given *r*⋆ using the Beverton-Holt model.

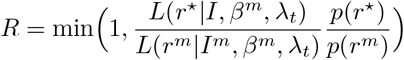

where *p* is the prior distribution of *r*. We define *p* ~ Gamma(*a_r_*, *b_r_*).

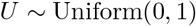 If *U* < *R* then *r*^*m*+1^ = *r*⋆. Else, *r*^*m*+1^ = *r^m^*.
6. Update *β*^*m*+1^. Propose *β*⋆ ~ Normal(*β*^*m*^, 0.2). The proposal is constrained to (0,1).

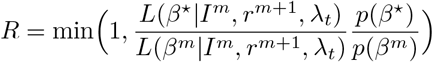

where *p* is the prior distribution of *β*. We define *p* ~ Beta(*α_β_*, *b_β_*).

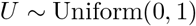 If *U* < *R* then *β*^*m*+1^ = *β*⋆. Else, *β*^*m*+1^ = *β*^*m*^.
7. Propose moving, adding or removing an infestation, each with equal probability. To move an infestation:

a. Update I. Select uniformly from the set of infested houses, *N_I_*. If *i* has not yet been inspected, propose new infestation time 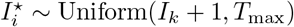. If *i* has been inspected but no insects were found at that time, propose new infestation time 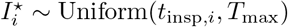. If *i* has been inspected and at least one insects was found at that time, propose new infestation time 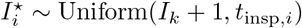.

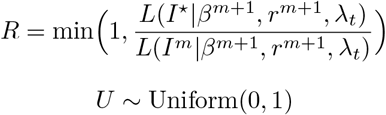 If *U* < *R* then *I*^*m*+1^ = *I*⋆. Else, *I*^*m*+1^ = *I^m^*. If *I*^*m*+1^ = *I*⋆, update *b_i_* (all insect counts corresponding to this infestation) so that at *I_i_*, *b*_*i*1_ = 1. To add an infestation:

a. Propose *i* uniformly from 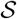, the set of susceptible houses.
b. If *i* has not yet been inspected, propose 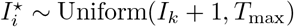. If *i* has been inspected, propose 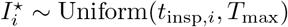. Propose 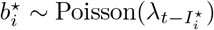.

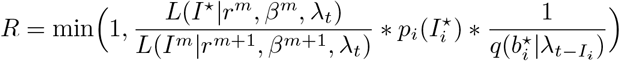

where *p_i_* is the prior probability that house *i* is infested and *q* is the proposal distribution of the insect counts *b_i_* (Poisson).

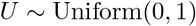 If *U* < *R* then *I*^*m*+1^ = *I*⋆. Else, *I*^*m*+1^ = *I^m^*. If a house is added, the corresponding insect counts but also be added. To remove an infestation:

a. Propose *i* uniformly from the set of occult infestations (previously added, but unobserved, houses).

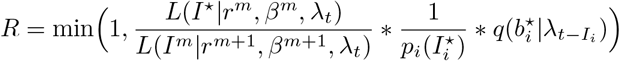

where *p_i_* is the prior probability that house *i* is uninfested and *q* is the distribution of the insect counts *b_i_* (Poisson).

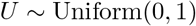 If *U* < *R* then *I*^*m*+1^ = *I*⋆. insect counts corresponding with the removed infestation must also be deleted. Else, *I*^*m*+1^ = *I^m^*.
8. Repeat steps (e) - (g) for a total of M iterations.

In the field implementation, *r* was not estimated, and thus step 5 was omitted. In general, the acceptance rates of parameters ranged from 15% to 60%, including those in the reversible-jump component. Posterior probabilities of infestation ranged from 0% to 20%, but mostly stayed below 10%.

Parameter estimates converged quickly, but it was difficult to assess convergence of posterior probabilities of infestation. We visually assessed convergence by plotting the ranking of each house between chains of the RJMCMC (Fig 5). Rankings were consistent in that top ranked houses were ranked highly across all chains. However, the specific ranking of a given house varied between chains. A typical convergence plot is shown below. A few houses seemed to get stuck at high rankings each chain that did get picked up in other chains. By using the median ranking for each house across chains, we hope to minimize this effect on inspections.

The simulation code is available at https://github.com/ebillig/Search-Strategy.

### Additional details

For all simulations and data applications, we fixed the carrying capacity, *K* = 1000. We did some sensitivity analysis to assess the importance of this assumption. We ran 3 RJMCMC chains on one locality with 5 different carrying capacities, *K* = {100, 500, 800,1000,1500}. We obtained the median ranking of each house across the chains for each *K*, and then plotted the rankings against each other (Fig 6). We can see the heterogeneity in ranking is similar to that between chains within each carrying capacity. Interestingly, the rankings stayed the same between *K* = 1000 and *K* = 1500.

